# AEMB: efficient abundance estimation for metagenomic binning

**DOI:** 10.1101/2025.07.30.667338

**Authors:** Shaojun Pan, Ivan Tolstoganov, Kristoffer Sahlin, Marcel Martin, Xing-Ming Zhao, Luis Pedro Coelho

## Abstract

Metagenomic binning is a crucial step in metagenomic analysis, namely grouping together contigs that are predicted to originate from the same genome to enable the recovery of metagenome-assembled genomes (MAGs). It has been shown that using information from multiple samples yields better results than binning each sample independently. However, for *N* metagenomic samples, using full multi-against-multi binning requires *N*^2^ alignments, making it computationally challenging to apply in large-scale metagenomic studies.

Here, we propose AEMB (Abundance Estimation for Metagenomic Binning), a novel mapping mode implemented in strobealign. AEMB is a computationally efficient abundance estimation method that uses a prefix-lookup vector as an indexing structure to reduce memory usage and randstrobes to estimate the abundance of contigs without performing base-level alignment. Compared to the hash table used in the previous version of strobealign, the indexing structure reduces peak memory usage by 25.2% with almost the same runtime. Furthermore, we implemented a fast abundance estimation method that skips base-level alignment. Altogether, AEMB reduces the runtime for abundance estimation by 88% to 96% compared to commonly used alignment methods such as Bowtie2 and BWA, while achieving similar binning results.

AEMB is available as a mapping mode in strobealign https://github.com/ksahlin/strobealign and SemiBin2 (v2.1 and later) accepts its inputs for binning.

## 1 Introduction

Metagenomic sequencing technology, commonly used for studying microorganisms that are difficult to culture^1^, has led to the generation of extensive collections of metagenome-assembled genomes (MAGs)^2,3^. This development significantly broadened our understanding of bacterial diversity across environments^4–7^, and elucidated the connections between microbiome composition and human health^8^.

*Metagenomic binning*, namely grouping contigs that are predicted to originate from the same genome, is a pivotal step in the processing of metagenomic data. Most widely-used metagenomic binning methods are reference-independent, relying on tetramer frequencies and abundance information, and include tools such as SolidBin^9^, MetaBAT2^10^, MaxBin2^11^, VAMB^12^, MetaDecoder^13^ and SemiBin1^14^ for short-read sequencing; and GraphMB^15^, LRBinner^16,17^, and SemiBin2^18^ for long-read sequencing. These methods perform binning by placing contigs with similar abundance into the same bin, with the assumption that these contigs originate from the same strain. The typical way to estimate contig abundances involves first aligning reads to contigs using tools such as Bowtie2^19^ or BWA^20^. Subsequently, the abundance for each contig is estimated from the resulting alignment file.

Often multiple related samples are available and different assembly and binning strategies can be employed (see Suppl. Fig. 1 and Methods). Single-sample binning, whereby each sample is treated independently, is commonly used in large-scale metagenomic analysis due to its computational efficiency, as it requires only a single read-alignment job between each sample’s reads and its resulting contig assembly^2,5,21–25^. Compared to single-sample binning, using abundance information from multiple samples yields better results, avoiding chimeric bins that may go undetected with single-sample binning^26^. However, the most straightforward way of calculating the abundance from *N* assemblies on *N* samples requires *N*^2^ alignment jobs, which has limited its applicability in large studies. An alternative approach aligns reads from multiple samples to a reference containing the contigs from all samples concatenated together, only requiring *N* alignments. However, this mode uses more time for the alignment of each sample (as reads are mapped to a larger reference) and may result in inaccurate estimates of abundance as there will be contigs from multiple samples that are very similar (even identical). This mode was introduced as *multisplit* binning in VAMB^12^ and refered to as *multi-sample binning* in SemiBin^14^ and SemiBin2^18^. For clarity, we will refer to this mode as *contacatenated-sample* binning.

Strobealign^27^, a recently proposed read alignment method, is much faster than traditional aligners, taking advantage of a novel seed type (based on syncmers^28^ and randstrobes^29^) and the fast query speed of a hash table. Despite its increased speed, strobealign was shown to have a substantially higher memory usage than tools based on BWT-FM^27,30^. Furthermore, strobealign’s full alignment mode outputs individual read alignments, which results in large output files and extra computational time in processing them.

To make the multi-against-multi approach viable for large datasets, we propose AEMB, a computationally efficient abundance estimation method for metagenomic binning (Fig. 1). To address the memory issue, we introduce an indexing structure based on a prefix-lookup vector as a substitute for the hash table used in previous versions of strobealign. This alternative reduces memory consumption without compromising runtime. Prefix-lookup vectors have been shown to be an effective alternative to hash tables in other scenarios, such as searching a suffix array for spliced alignment of RNA-seq reads^31^. For abundance estimation, AEMB uses randstrobes^29^ to match reads and contigs and estimate the abundance of contigs without performing base-level alignment. On simulated and real data, we show that the abundance estimates obtained by AEMB for binning yield similar results to commonly used alignment methods but significantly reduces runtime (88%-96%), enabling multi-against-multi binning for large metagenomic studies.

**Fig 1.**
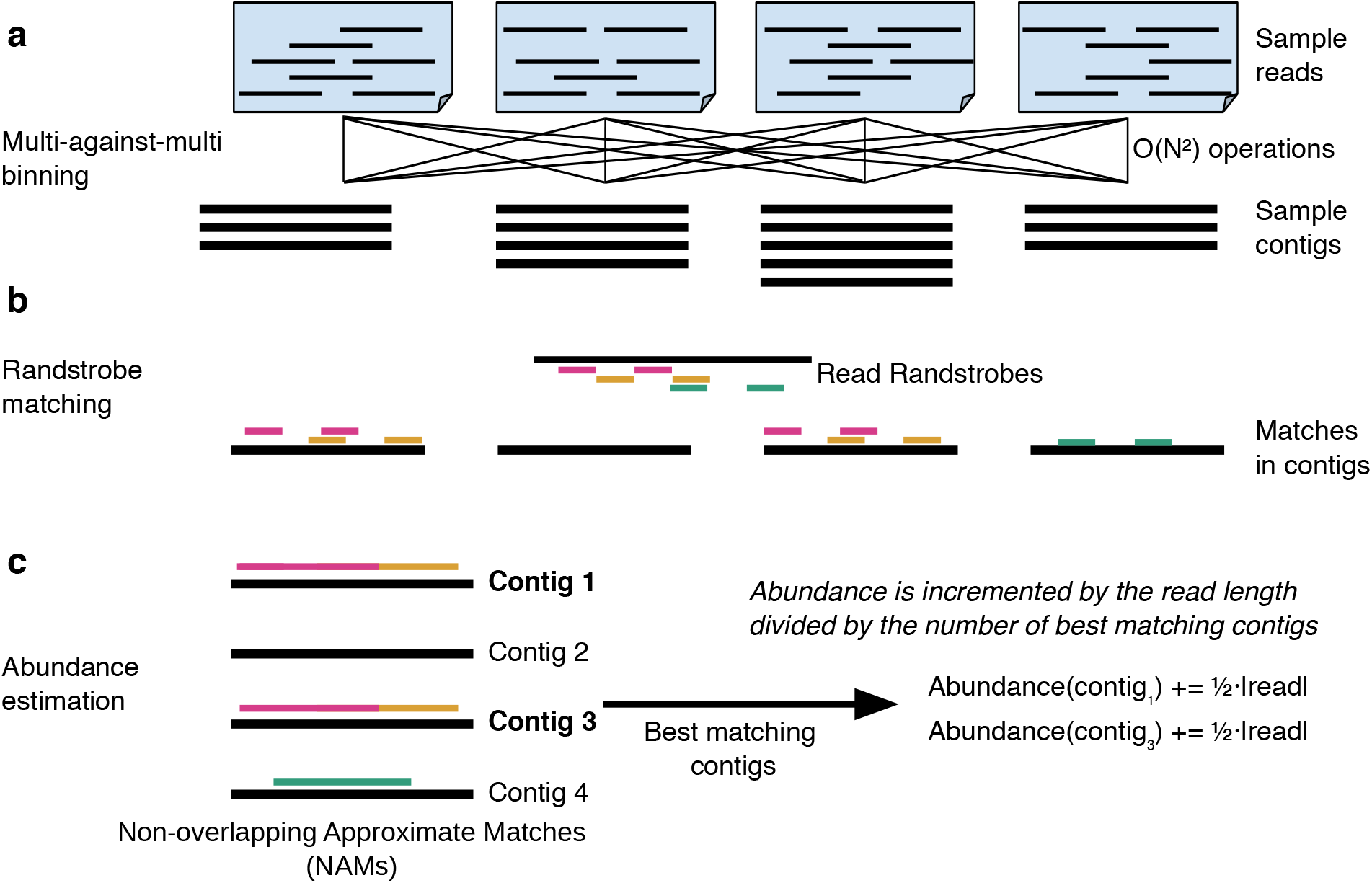
Overview of multi-against-multi mapping and the AEMB pipeline. **(a)** Multi-against-multi mapping, where each sample is mapped against every set of assembled contigs, results in *O*(*N*^2^) mapping jobs for *N* samples. **(b)** Randstrobes^27^ are constructed from the reads. The randstrobes with the same hash values between the read and the references are identified. **(c)** The randstrobes of a read are used to query AEMB’s indexing structure, and NAMs are constructed from the randstrobes to compute the matching score between the read and the contigs. Based on the best matching score, the read is assigned to the corresponding contig. In cases where multiple contigs share the same optimal matching score, the read is evenly distributed among these contigs.

## 2. Results and Discussion

### 2.1 The AEMB algorithm

Strobealign uses long approximate seeds, which have been demonstrated to considerably reduce seed repetitiveness compared to other methods^27^. The reduced repetitiveness of the seeds, together with a hash-table-based index implementation, enables much faster mapping than other read mappers at comparable accuracy^27^. However, strobealign was shown to have a substantially higher memory usage than other tools^27^. Over 50% of the memory use stems from the hash table index. In addition, it outputs detailed information of each read mapping, which is not necessary for abundance estimation.

We developed AEMB, a computationally efficient abundance estimation method for metagenomic binning (see Fig. 1). The AEMB algorithm takes advantage of strobealign’s seeding method but improves upon its hash-table-based indexing structure (see Methods). It also enables fast and accurate abundance estimate computations.

The main insight comes from the fact that the index is static during mapping. After its construction, we do not need many of the functionalities that a hash table offers, such as deleting and reinserting elements. This enables a more memory-efficient look-up structure based on a prefix array (details in Methods). After indexing, AEMB calculates the matching scores between each input read and contigs (see Methods) by querying the index. AEMB assigns the read to the contig with the highest score. If multiple contigs have the same matching score, AEMB distributes the read evenly among them. Finally, AEMB multiplies the allocated number of reads by the length of the reads and divides it by the length of the contig to obtain the abundance of the contig (see Methods).

### 2.2 The indexing structure consumes less memory than a hash table

We compared the performance of our proposed indexing structure with the hash table used in previous versions of strobealign. We utilized a soil dataset^32^, aligning ten randomly selected samples to one contig file (one indexing, ten mappings). For the comparison with strobealign’s hash table implementation, we directly replaced the indexing structure in AEMB with strobealign’s hash table (see Methods), which we denote as AEMB(hash).

In AEMB, *B* is a parameter controlling the size of the indexing structure (details in Methods). The prefix array is of size 2^*B*^, so memory usage grows with *B* On the other hand, runtime decreases with increasing *B*, in a classical memory/runtime tradeoff (see Fig. 2). AEMB estimates the optimal value of *B* based on the number of seeds, denoted AEMB(default); see Methods. Users can also adjust the *B* based on their specific needs (if they have more memory available, they can set a larger *B* to reduce runtime). As shown in Fig. 2, AEMB(default), which sets *B* = 27 in this dataset, incurred a slight increase in runtime compared to the hash table index (12.4% increase per sample) but reduced the peak memory usage by 40.2%. Setting *B* = 29 leads to nearly identical runtime (1.7% slower) to the hash-based index, while still reducing the memory usage by 25.7%.

**Fig 2.**
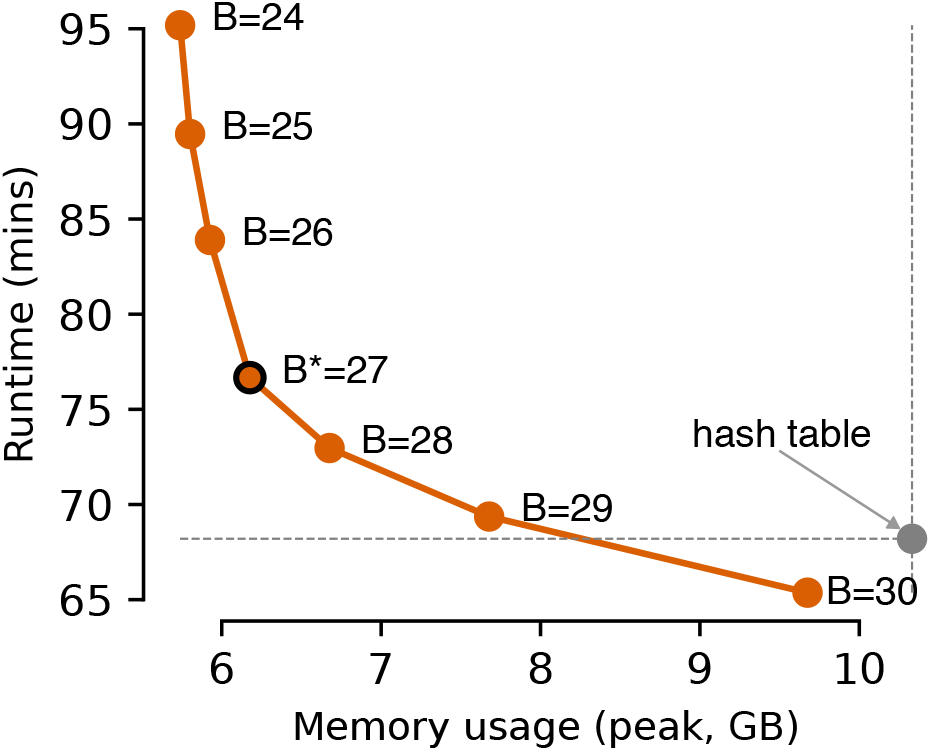
The indexing structure requires less peak memory and similar runtime compared to hash table. Runtime and peak memory usage of AEMB with different values of *B* and the hash-table AEMB in the soil environment. In this dataset, the heuristic method chooses *B* = 27 as default.

We compared the number of high-quality distinct taxa produced by SemiBin2^18^ using abundance features from AEMB(hash) and AEMB(default). Five simulated datasets from the CAMI II challenge^33,34^ were used for the benchmark (see Methods). In the five environments, AEMB(hash) and AEMB(default) exhibited similar results based on the multi-against-multi binning mode (see Suppl. Fig. 2, Methods). The slight difference may be explained by the fact that the sequence matches used to estimate abundance depend on the seed order in the index^27^, which differs between AEMB(hash) and AEMB(default) (see Methods).

### 2.3 AEMB is much faster compared to alignment-based abundance estimates

We compared binning results obtained using AEMB-produced abundance features with those from alignment-based methods. We used Bowtie2^19^ and BWA^20^, two widely used alignment tools, as well as the full alignment-based version of strobealign^27^. In the CAMI II simulated datasets, multi-against-multi binning (see Methods) with abundance information from AEMB and base-level alignment methods reconstructed similar numbers of distinct high-quality genera, species and strains (see Fig. 3).

**Fig 3.**
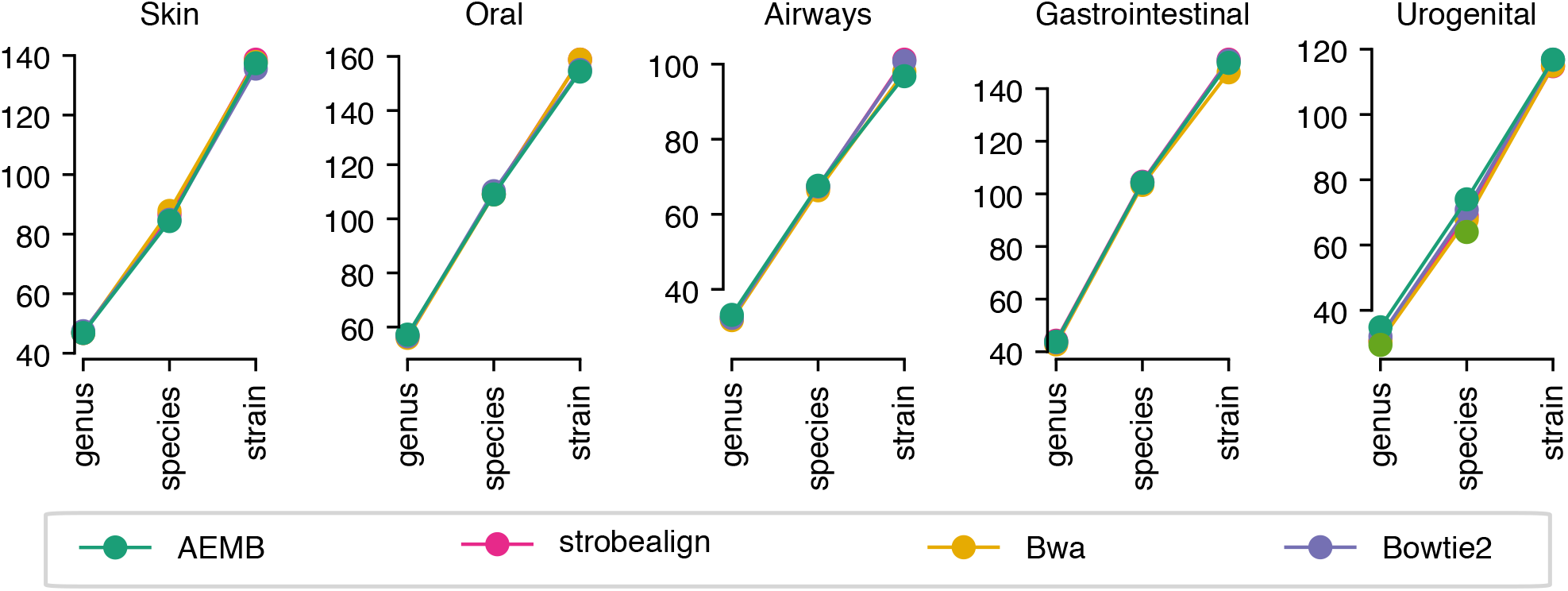
AEMB and alignment-based methods performs similarly in CAMI II datasets when used for binning. Shown are the binning results with multi-against-multi mode (the number of distinct genera, species and strains) with the abundance information from AEMB and alignment-based abundance estimation in the CAMI II datasets.

In every environment, we performed indexing only once and then aligned reads from all samples to the contigs. AEMB resulted in significantly reduced runtime compared to other methods. Compared to Bowtie2 and BWA, AEMB reduced the average runtime by 96.1% and 95.7%, respectively (see Fig. 4). When compared to the classical version of strobealign, it still reduced the runtime by 88.5% (see Fig. 4). These results indicate that AEMB significantly reduces the time for abundance calculations without affecting binning results, enabling multi-against-multi binning mode in large-scale metagenomic analyses.

**Fig 4.**
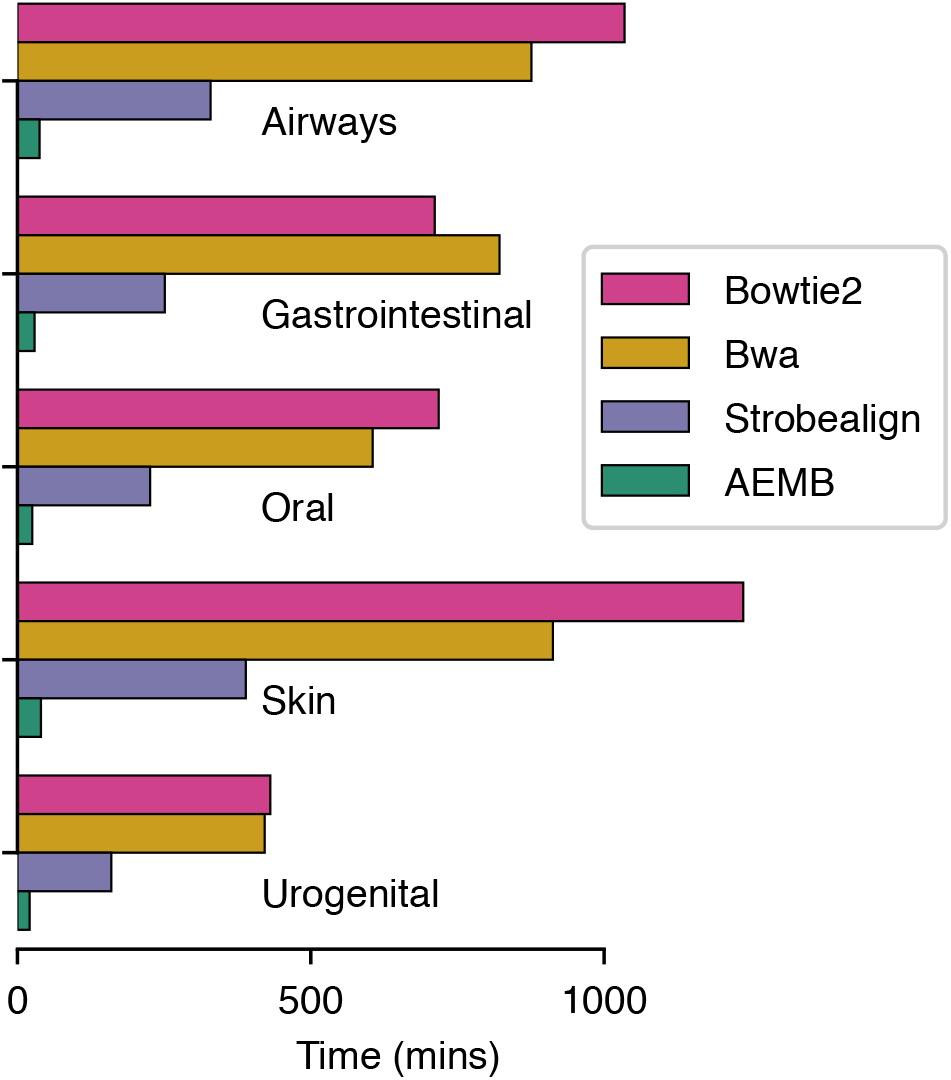
AEMB is much faster than other alignment-based abundance calculation methods. Runtime of AEMB and alignment-based approaches (Bowtie2, BWA, strobealign) to estimate abundances in CAMI II datasets.

### 2.4 AEMB speeds up binning in real data

To evaluate AEMB on real data, we used four different environments: human gut (n = 82), dog gut (n = 129), ocean (n = 109) and soil (n = 101)^35–38^. For every environment, we randomly selected 20 samples as testing data and calculated abundance features using AEMB on the testing data with different sample sizes (see Methods). Subsequently, binning was performed with SemiBin2. We used CheckM^39^ and GUNC^40^ to evaluate the bins (see Methods).

We compared the binning results obtained from abundance features calculated using AEMB and Bowtie2. We compared AEMB to Bowtie2 with single-sample (Bowtie2(single), using a pretrained model in SemiBin1^14^) and concatenated-sample (all samples used in the environment, Bowtie2(concat)) binning mode. It is impractical to use Bowtie2 for multi-against-multi binning on such a large-scale dataset.

We observed that when a larger number of samples is used, AEMB yields results similar to, or even better than Bowtie2 with concatenated-sample binning (see Fig. 5). In the human gut environment, when ≥ 30 samples are used, AEMB can reconstruct an average of 11.1% (6.7%-15.4%) more high-quality bins than Bowtie2(concat) using 82 samples and in the dog gut environment, when the number of samples used ≥ 40, AEMB can reconstruct an average of 4.0% (1.9%-5.6%) more high-quality bins compared to Bowtie2(concat) using 129 samples. In the soil environment, when the number of samples used ≥ 50, AEMB achieves similar results as Bowtie2(concat) using 101 samples. In the ocean environment, AEMB exhibits a decrease (11.3%) compared to Bowtie2(concat) when using all 109 samples in the number of high-quality bins. However, considering the improvement in computational performance, the decrease in results in this environment is within an acceptable range.

**Fig 5.**
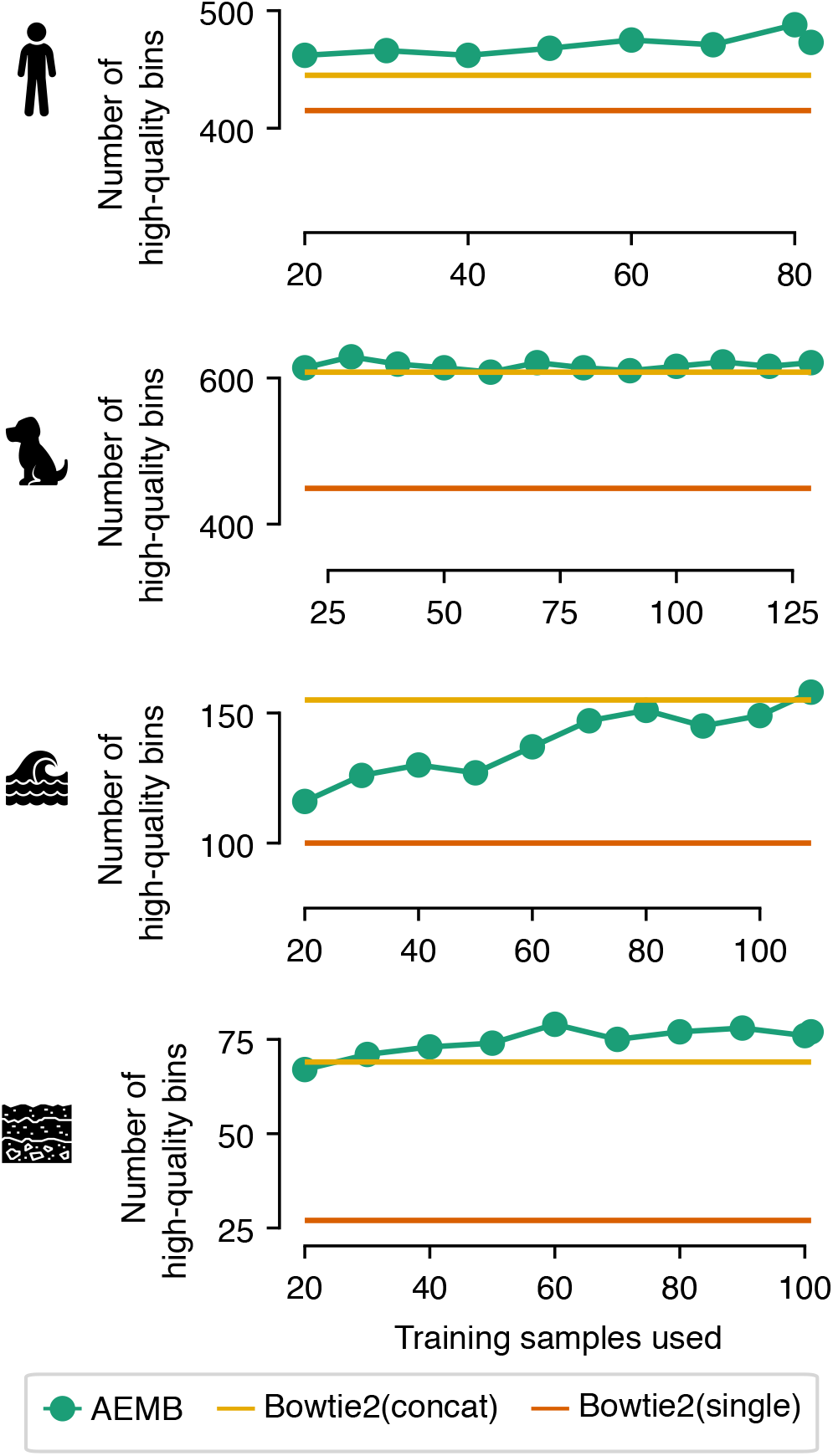
More samples can lead to better binning results in real data. For every environment (human gut, dog gut, ocean and soil), we sampled 20 samples as testing sets and used different numbers of samples to obtain abundances. Shown are the numbers of high-quality bins with different numbers of samples used in binning (abundance estimated with AEMB) and the number of high-quality bins with abundance information calculated from Bowtie2. Bowtie2(single) shows the result of single-sample mode, while Bowtie2(concat) shows the result of running SemiBin2 with the heuristic concatenated-sample binning.

In the four environments, Bowtie2(single) outperforms Bowtie2(concat) only in the human gut dataset. The possible reason is that concatenated-sample binning aligns reads to the concatenated contigs, leading to less accurate abundance estimation. When using AEMB results, AEMB outperforms Bowtie2 (single) when the number of samples is 30 or more.

As more samples are used for binning, the number of resulting high-quality bins initially rises and then the growth tends to plateau, in an environment-dependent manner (see Fig. 5). Therefore, beyond a certain number, using more samples for binning does not necessarily lead to better results, but also does not degrade the results. The optimal choice of the number of samples for binning should be considered based on the specific circumstances, balancing the computational cost and quality of the results.

## 3 Conclusions

Large-scale genome recovery from microbiomes will remain relevant for the foreseeable future, as even with recent efforts, there are still many microbial species without a genome available in public databases^41–43^. Despite widespread agreement that single-sample binning does not return the best results, the high computational cost of multi-against-multi binning has limited its use in large-scale metagenomic analyses. Other studies approximated multi-against-multi binning by combining all the contigs into a single database and aligning the reads to this single combined database, reducing the number of alignment jobs from *N*^2^ to *N*, where *N* is the number of samples^12,14,18^. This approach is valuable and available in SemiBin2^18^, but it may not accurately estimate the abundance of each individual contig, particularly when contigs from different samples are very similar or even identical. Empirically, this approach showed a large improvement compared to single-sample binning, but it was still not as good as multi-against-multi binning (see Fig. 5).

Tools like BWA, Bowtie2, and the full-alignment mapping mode implemented in strobealign, use base-level alignment to assign reads to contigs, which is accurate but computationally expensive. We developed the AEMB abundance estimator which circumvents base-level alignment by assigning reads to contigs based on matching scores from randstrobe seeds. When used for binning in multi-against-multi mode, AEMB abundance estimations result in a similar number of high-quality bins to base-level alignment. However, compared to state-of-the-art alignment methods, AEMB reduces the time needed to calculate abundance by 88% to 96%, and as such can be used in large-scale metagenomic analysis.

## 4 Methods

We first describe strobealign’s indexing layout in order to discuss the differences introduced by AEMB’s approach.

### 4.1 Strobealign’s index layout

Strobealign requires mapping from a hash representation of a seed to its position in the reference. For each seed, strobealign stores a hash value representation of the seed (64 bits), the seed length (8 bits), and the seed location (sequence ID and position; 56 bits) in a flat vector sorted based on the seed hash values. Each entry in the flat vector therefore contains a 128-bit entry. Since the vector is sorted by seed hash value, identical seeds appear next to each other (e.g., if multiple contigs share the same region). From now on, we refer to the sorted flat vector with seed information as *R*. Previously, strobealign also included a hash table that associates the seed hash value with the seed location in the flat vector (32 bits) and its count (repetitiveness; 32 bits) as the value. Therefore, each entry in the hash table also takes 128 bits.

Since the large majority of seeds in strobealign are unique for typical genomes ^27^, this makes the hash table use nearly as much memory as *R*. In addition, the hash table needs additional slots to keep a low load factor, for efficient insertion of future elements.

### 4.2 AEMB pipeline

#### 4.2.1 AEMB’s indexing structure

The main insight comes from the observation that after its construction, *R* is static and we do not need to delete or insert new elements, which would require keeping a low load factor. Here, we introduce a prefix-lookup-based indexing structure to replace the commonly used hash table in hash-based alignment algorithms.

We use a prefix-lookup vector denoted by *S*. The vector *S* is of size 2^*B*^ (*B* is a parameter), where each entry is a 64-bit integer. Let *S*_*n*_ denote the integer at position *n* in *S*. Then, *S*_*n*_ stores the first position in *R* such that the top *B* bits (i.e., the most significant *B* bits) of its seed hash value is *n* (see Fig. 6). For example, if *B* = 8 and *S*_4_ = 137 (recalling that 4=00000100 in binary form with *B* = 8), this means that *R*_137_ contains a hash value starting with 00000100. Furthermore, if *S*_5_ = 138, it implies that the seed with prefix 00000100 is unique in the index. A larger *B* gives more accurate position approximations, but requires more memory.

**Fig 6.**
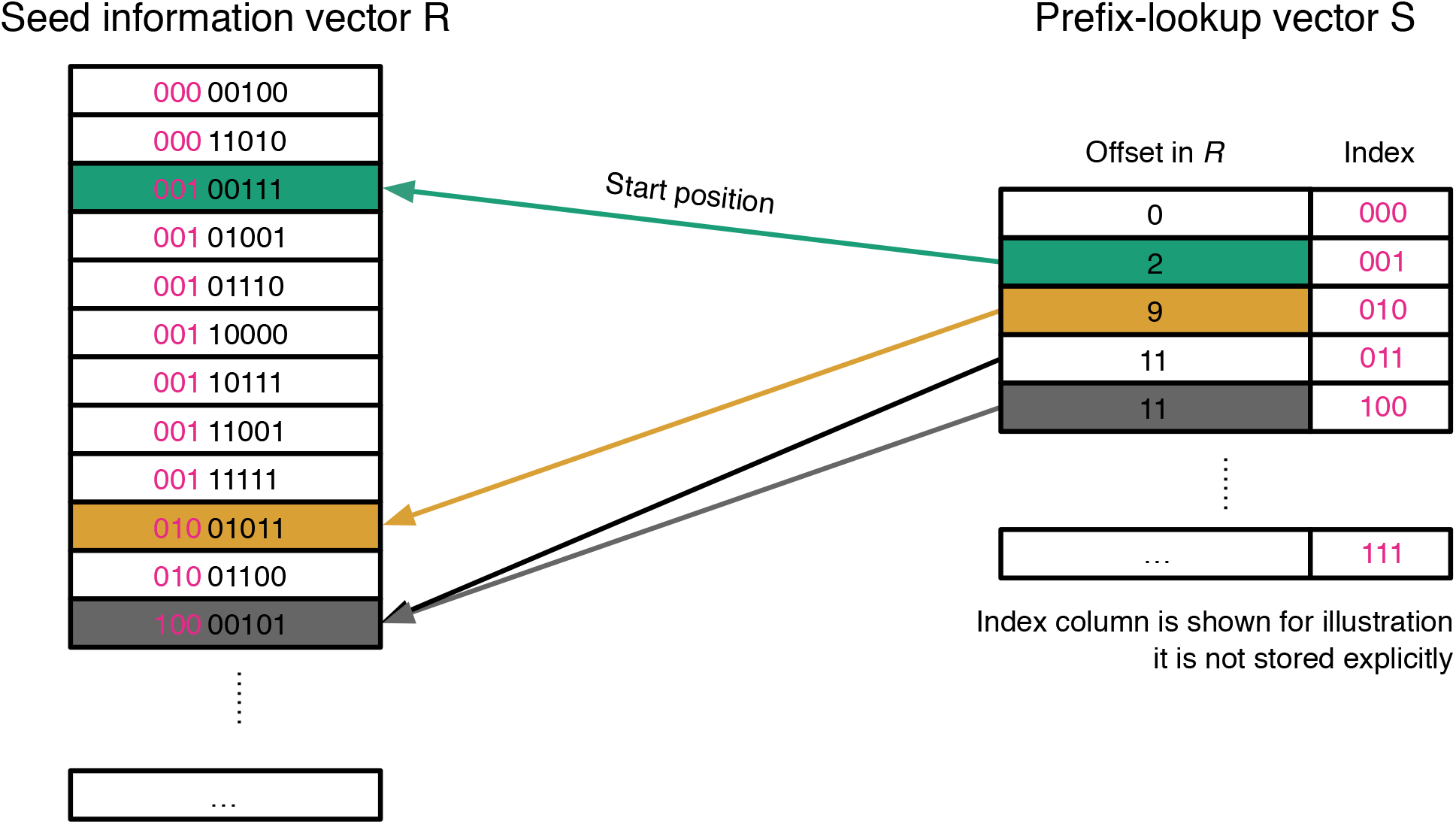
The AEMB index structure. The indexing layout of strobealign consists of two structures, the sorted flat vector *R* of the seed hash values, sequence ID, and positions (left), and a prefix-lookup vector (right). At the position *n*, the prefix-lookup vector stores the first position in *R* with *n* as the prefix of size *B*. If there is no such position in *R, S*_*n*_ points instead to the next available starting position.

#### 4.2.2 Constructing the prefix-lookup vector

Let a seed in the index vector *R* have the hash value *h* and be positioned at index *p* in *R* (so that *R*_*p*_ = *h*), and the top *B* bits of the hash represented as *T*_*B*_(*p*). Since *R* is sorted with respect to *h*, it is also sorted with respect to *T*_*B*_(*p*). Thus, elements with the same values *T*_*B*_(*p*) appear consecutively in *R*. We denote the set of all possible top *B* bits of hashes in *R* as *T*_*B*_(*R*). For every *t* ∈ [0, 2^*B*^ − 1], we define the value at position *t* in the prefix lookup vector *S* as follows:

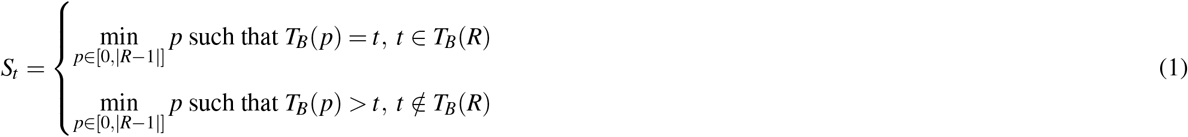

The second case in (1) corresponds to the situation when *t* is not encountered as the top *B* bits of any hash value in *R*. In that case, we assign the position of the next existing hash value prefix to *S*_*t*_. That is, assume *t, t* + 1, …, *t* + *d* − 1 do not exist in the top *B* bits in *R* but *t* + *d* exists. Then positions *t, t* + 1, …, *t* + *d* − 1 in *S* will store the position of *t* + *d* in *R*. With this construction, it becomes easy to obtain the starting and ending positions of the hash prefix *t* in *R* through *S*_*t*_ and *S*_*t*+1_. However, if *t* + 1 does not exist in *T*_*B*_(*R*), we still find the end position of *t* as *t* + *d* is the next occurring hash value prefix in *R*.

The rules above allow an efficient streaming procedure when building the index. The process of constructing the full index is as follows:

1. Construct seeds from the reference sequences using the same approach as in strobealign^27^.
2. Store the hash values and position information of each seed in the vector *R*, and sort them in-place in ascending order based on the hash values.
3. Allocate the prefix-lookup vector *S* with size 2^*B*^ and update each position of this vector according to the method mentioned above. Specifically, iterate over the elements in *R* and update each position *t* ∈ [0, 2^*B*^ − 1] in *S* in ascending order according to (1).

When saving the index to disk, we store the prefix-lookup vector *S* and the vector *R* directly in binary format, which makes loading the index into memory very fast.

#### 4.2.3 Querying the prefix-lookup vector

During the querying process, firstly, AEMB constructs seeds from the input read and then utilizes the top *B* bits of the hash value from the queried seeds to search *R*. Specifically, as above, if the hash value of the query seed is *h* and the top *B* bits of *h* is *t*, then we query the two positions, *S*_*t*_ and *S*_*t*+1_. By construction, *S*_*t*_ ≤ *S*_*t*+1_. If *S*_*t*+1_ − *S*_*t*_ *<* 4, then a simple linear search is used, checking all positions in the interval [*S*_*t*_, *S*_*t*+1_). Note that, as a special case, if *S*_*t*_ = *S*_*t*+1_, no entries in *R* need to be checked and the process returns early. Otherwise, we start a binary search in *R* using *S*_*t*_ and *S*_*t*+1_ as initial start and end points.

*B* can be set by the user, but by default, a heuristic is used, based on the estimated number of randstrobes, *T*, namely *B* = max(8, ⌊log_2_ *T*⌋ − 1). Here, 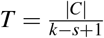, where |*C*| denotes the total length of all contigs and 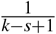 is the expected syncmer sampling density based on syncmer parameters *k* and *s*^27^, which determines the number of randstrobes.

#### 4.2.4 Abundance estimation

Randstrobes are extracted from each input read using the same parameters as were used during index construction. The index (previously a hash table, now the prefix vector described above) is then queried to find candidate matches between the read and the reference^27^. These matches are merged to Non-overlapping Approximate Matches (NAMs)^27,29^. AEMB uses the NAM matching score, *m*, to rank candidate contigs:

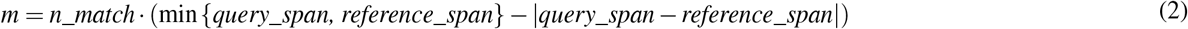

where *n*_*match* is the number of matches, *query*_*span* and *re f erence*_*span* are the length of merged matching in query and reference, respectively.

Based on the matching score *m*, AEMB finds the best matching contig for every input read. In the case of paired-end reads, each read mate is first processed independently and, if mates are assigned to different contigs, the match is split between them. If one read has multiple best matches (with the same matching score), AEMB evenly distributes the abundance of this read among multiple contigs. Finally, the estimated coverage of each contig is obtained by dividing by the length of the contig.

#### 4.2.5 Binning with SemiBin2

SemiBin2 needs abundance information for split contigs^14,18^ and AEMB cannot give the per-base coverage of reads mapped to the contig to calculate the abundance of split contig (which base-level alignment methods can give). To run SemiBin2 with AEMB, we divided the original contig into two shorter segments and used AEMB to align reads to the shorter contigs, obtaining abundance values for the separated contigs. We then calculated the mean of the abundances of the two shorter segments to obtain the abundance of the original contig.

#### 4.2.6 Binning modes

Multiple strategies are available for dealing with multiple samples (see Suppl. Fig. 1):

1. Single-sample binning: every sample is assembled and binned independently.
2. Co-assembly binning: all samples are co-assembled first, then binning is performed with abundance information from several samples. Co-assembly binning can use abundance information from several samples and recover lower abundance genomes^42^, but may result in chimeric contigs and, by design, cannot reconstruct sample-specific variation (e.g., different strains of the same species).
3. Multi-against-multi binning: every sample is assembled independently and reads from each sample are mapped to the contigs of every sample. Multi-against-multi binning can reconstruct sample-specific variation and use abundance information across samples. However, multi-against-multi binning requires *N*^2^ alignments, which limits its scalability.
4. Multisample or concatenated sample binning: every sample is assembled independently and all assemblies are combined to a concatenated file. Then all samples are mapped to the concatenated file to generate abundance information across all samples for every contig used in binning. This approach only requires *N* alignments, but aligning to a combined reference needs more time and the abundance estimation will be less precise.

### 4.3 Data used

For benchmarking, we used five simulated datasets from the CAMI II challenges: airways (10 samples), gastrointestinal (10 samples), oral (10 samples), skin (10 samples), urogenital (9 samples), and four real datasets: human gut (82 samples)^35^, dog gut (129 samples)^36^, ocean (109 samples)^37^ and soil (101 samples)^32^.

### 4.4 Methods used in benchmarking

To run AEMB with the hash table version, we used the hash table implementation in the previous version of strobealign^27^, with the abundance estimation method described in this paper.

When calculating abundance with Bowtie2 (version 2.5.2), BWA (version 0.7.17), and strobealign (version 0.12.0), reads are first aligned to contigs using these tools. All tools were run with four threads. Subsequently, abundance is computed using BEDTools^44^ (version 2.31.1, using the `genomecov` subcommand; this is implemented internally in SemiBin2). When evaluating the runtime and peak memory usage, we only performed the indexing once.

To benchmark AEMB in real data, we used four different environments: human gut, dog gut, ocean, and soil. We randomly sampled 20 samples as the testing set. Subsequently, we randomly selected 20, 30, 40, 50, 60, 70, 80, 82 samples for human gut; 20, 30, 40, 50, 60, 70, 80, 90, 100, 110, 120, 129 samples for dog gut; 20, 30, 40, 50, 60, 70, 80,90,100,109 samples for ocean and 20, 30, 40, 50, 60, 70, 80,90,100,101 samples for soil environment as abundance features for binning (aligning these samples using AEMB to the testing data to get abundance features).

### 4.5 Evaluation metrics

For the evaluation of binning, we used completeness and contamination to evaluate a bin. For the simulated datasets, there are gold standard assemblies from CAMI II challenge and we can use AMBER^45^ (version 2.0.2) to calculate the completeness and contamination of every bin. For real datasets, given that there is no known ground truth, we used CheckM^39^ (version 1.1.9) and GUNC^40^ (version 1.0.5) to evaluate the bins.

For the simulated datasets, a high-quality bin is defined as having completeness > 90% and contamination < 5%. For the biological datasets, a high-quality bin is defined as having completeness > 90%, contamination < 5% and passing GUNC’s chimera detection. The number of genera, species, and strains having at least one high-quality bin associated with them is reported as the number of distinct genera, species, and strains, respectively.

## 4.6 Code availability

AEMB is available as a mapping mode in strobealign https://github.com/ksahlin/strobealign since version 0.13. Binning from AEMB’s abundance estimates is available in SemiBin2 since version 2.1 and SemiBin2’s source code is available at https://github.com/BigDataBiology/SemiBin. The analysis script used in the study to benchmark the tools can be found at https://github.com/BigDataBiology/AEMB_benchmark.

## 4.7 Acknowledgements

This work was supported by the SciLifeLab & Wallenberg Data Driven Life Science Program, Knut and Alice Wallenberg Foundation (grants: KAW 2020.0239 and KAW 2017.0003), and by the National Bioinformatics Infrastructure Sweden (NBIS) at SciLifeLab. Kristoffer Sahlin was supported by the Swedish Research Council (SRC, Vetenskapsrådet) under Grant No. 2021-04000. Luis Pedro Coelho is supported by the Australian Research Council (grant FT230100724).

We thank the users of strobealign and SemiBin for their suggestions and bug reports.

## 4.8 Author Contributions

SP, KS, and LPC conceived and designed the algorithms. SP, KS, MM, and LPC implemented the algorithms. SP performed the experiments. KS, XM, and LPC supervised the project. SP, KS, IT, XM, and LPC wrote the manuscript and designed the figures. All authors contributed to the revision of the manuscript prior to submission.

## 6 Supplementary Figures and Tables

**Supplementary Fig 1.**
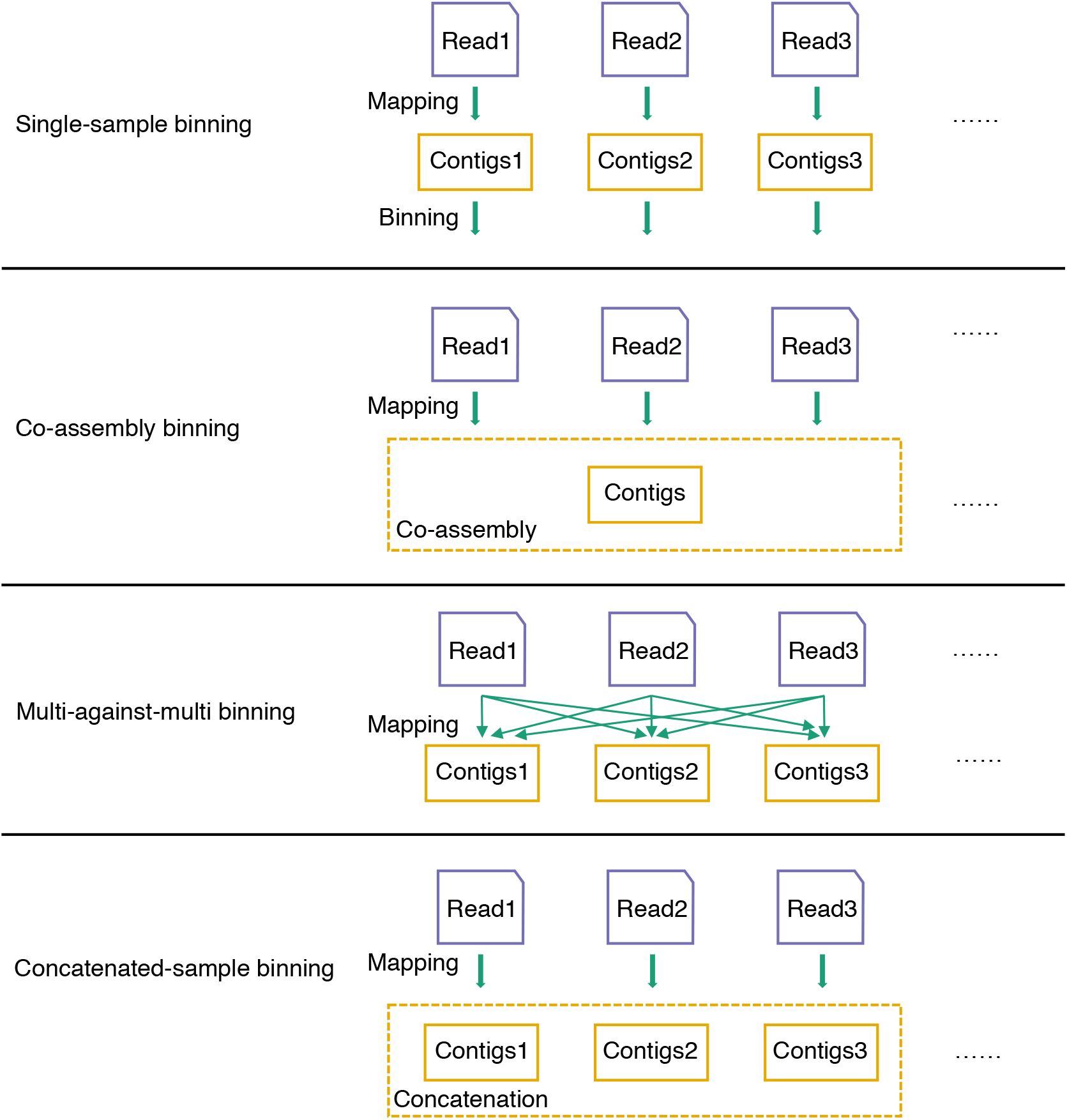
Different binning modes. Single-sample binning means every sample is assembled, mapped and binned independently. Co-assembly binning means all samples are co-assembled first and then binning with abundance information from several samples. Multi-against-multi binning means every sample is assembled independently and then reads from all samples are mapped to the contigs of every sample, then binning is performed for every sample with abundance information from several samples. Concatenate-sample binning also assembles every sample independently and combines these contigs together, then reads from every sample are mapped to a concatenated contig file to generate abundance information across several samples used in binning.

**Supplementary Fig 2.**
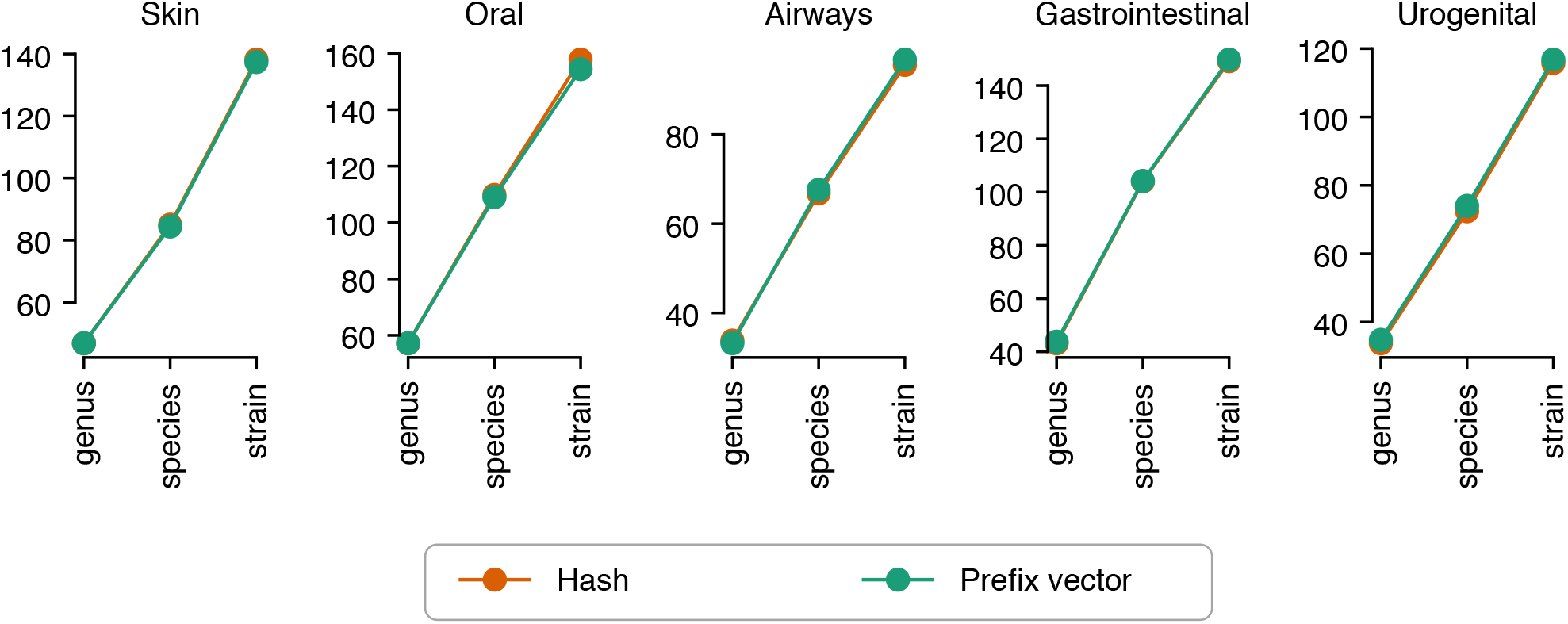
The indexing structure and hash table performs similarly in CAMI II datasets when used for binning. Shown are the binning results with multi-against-multi mode (the number of distinct genera, species and strains) with the abundance information from proposed indexing structure and hash table in the CAMI II datasets.

## References

1. Quince, C., Walker, A. W., Simpson, J. T., Loman, N. J. & Segata, N. Shotgun metagenomics, from sampling to analysis. Nat. Biotechnol. 35, 833–844 (2017).

2. Coelho, L. P. et al. Towards the biogeography of prokaryotic genes. Nature 601, 252–256 (2022).

3. Schmidt, T. S. B. et al. SPIRE: a searchable, planetary-scale mIcrobiome REsource. Nucleic Acids Res. 52, D777–D783 (2024).

4. Tully, B. J., Graham, E. D. & Heidelberg, J. F. The reconstruction of 2,631 draft metagenome-assembled genomes from the global oceans. Sci. Data 5, 1–8 (2018).

5. Almeida, A. et al. A new genomic blueprint of the human gut microbiota. Nature 568, 499–504 (2019).

6. Stewart, R. D. et al. Compendium of 4,941 rumen metagenome-assembled genomes for rumen microbiome biology and enzyme discovery. Nat. Biotechnol. 37, 953–961 (2019).

7. Zeng, S. et al. A compendium of 32,277 metagenome-assembled genomes and over 80 million genes from the early-life human gut microbiome. Nat. Commun. 13, 1–15 (2022).

8. Jaenicke, S., Diedrich, S. & Goesmann, A. Mgx 2.0: Shotgun-and assembly-based metagenome and metatranscriptome analysis from a single source. bioRxiv 2023–09 (2023).

9. Wang, Z., Wang, Z., Lu, Y. Y., Sun, F. & Zhu, S. SolidBin: improving metagenome binning with semi-supervised normalized cut. Bioinformatics 35, 4229–4238 (2019).

10. Kang, D. D. et al. MetaBAT 2: an adaptive binning algorithm for robust and efficient genome reconstruction from metagenome assemblies. PeerJ 7, e7359 (2019).

11. Wu, Y.-W., Simmons, B. A. & Singer, S. W. Maxbin 2.0: an automated binning algorithm to recover genomes from multiple metagenomic datasets. Bioinformatics 32, 605–607 (2016).

12. Nissen, J. N. et al. Improved metagenome binning and assembly using deep variational autoencoders. Nat. biotechnology 39, 555–560 (2021).

13. Liu, C.-C. et al. MetaDecoder: a novel method for clustering metagenomic contigs. Microbiome 10, 1–16 (2022).

14. Pan, S., Zhu, C.Zhao, X.-M. & Coelho, L. P. A deep siamese neural network improves metagenome-assembled genomes in microbiome datasets across different environments. Nat. Commun. 13, 1–12 (2022).

15. Lamurias, A., Sereika, M., Albertsen, M., Hose, K. & Nielsen, T. D. Metagenomic binning with assembly graph embeddings. Bioinformatics 38, 4481–4487 (2022).

16. Wickramarachchi, A., Mallawaarachchi, V., Rajan, V. & Lin, Y. MetaBCC-LR: metagenomics binning by coverage and composition for long reads. Bioinformatics 36, i3–i11 (2020).

17. Wickramarachchi, A. & Lin, Y. LRBinner: Binning Long Reads in Metagenomics Datasets. In 21st International Workshop on Algorithms in Bioinformatics (WABI 2021) (2021).

18. Pan, S.Zhao, X.-M. & Coelho, L. P. SemiBin2: self-supervised contrastive learning leads to better MAGs for short- and long-read sequencing. Bioinformatics 39, i21–i29, DOI: 10.1093/bioinformatics/btad209 (2023). https://academic.oup.com/bioinformatics/article-pdf/39/Supplement_1/i21/50741692/btad209.pdf.

19. Langmead, B. & Salzberg, S. L. Fast gapped-read alignment with Bowtie 2. Nat. Methods 9, 357–359 (2012).

20. Li, H. Aligning sequence reads, clone sequences and assembly contigs with bwa-mem. arXiv preprint 1303.3997 (2013).

21. Pasolli, E. et al. Extensive unexplored human microbiome diversity revealed by over 150,000 genomes from metagenomes spanning age, geography, and lifestyle. Cell 176, 649–662 (2019).

22. Nayfach, S., Shi, Z. J., Seshadri, R., Pollard, K. S. & Kyrpides, N. C. New insights from uncultivated genomes of the global human gut microbiome. Nature 568, 505–510 (2019).

23. Nayfach, S. et al. A genomic catalog of earth’s microbiomes. Nat. biotechnology 1–11 (2020).

24. Schulz, F. et al. Giant virus diversity and host interactions through global metagenomics. Nature 578, 432–436 (2020).

25. Asnicar, F. et al. Microbiome connections with host metabolism and habitual diet from 1,098 deeply phenotyped individuals. Nat. Medicine 27, 321–332 (2021).

26. Mattock, J. & Watson, M. A comparison of single-coverage and multi-coverage metagenomic binning reveals extensive hidden contamination. Nat. Methods 20, 1170–1173 (2023).

27. Sahlin, K. Strobealign: flexible seed size enables ultra-fast and accurate read alignment. Genome Biol. 23, 260 (2022).

28. Edgar, R. Syncmers are more sensitive than minimizers for selecting conserved k-mers in biological sequences. PeerJ 9, e10805 (2021).

29. Sahlin, K. Effective sequence similarity detection with strobemers. Genome research 31, 2080–2094 (2021).

30. Alser, M. et al. Technology dictates algorithms: recent developments in read alignment. Genome biology 22, 249 (2021).

31. Dobin, A. et al. STAR: ultrafast universal RNA-seq aligner. Bioinformatics 29, 15–21 (2012).

32. Olm, M. R. et al. The source and evolutionary history of a microbial contaminant identified through soil metagenomic analysis. MBio 8, e01969–16 (2017).

33. Sczyrba, A. et al. Critical assessment of metagenome interpretation—a benchmark of metagenomics software. Nat. methods 14, 1063–1071 (2017).

34. Meyer, F. et al. Critical assessment of metagenome interpretation: the second round of challenges. Nat. Methods 19, 429–440 (2022).

35. Wirbel, J. et al. Meta-analysis of fecal metagenomes reveals global microbial signatures that are specific for colorectal cancer. Nat. Med. 25, 679–689 (2019).

36. Coelho, L. P. et al. Similarity of the dog and human gut microbiomes in gene content and response to diet. Microbiome 6, 1–11 (2018).

37. Sunagawa, S. et al. Structure and function of the global ocean microbiome. Science 348, 1261359 (2015).

38. Olm, M. R. et al. The source and evolutionary history of a microbial contaminant identified through soil metagenomic analysis. MBio 8, e01969–16 (2017).

39. Parks, D. H., Imelfort, M., Skennerton, C. T., Hugenholtz, P. & Tyson, G. W. Checkm: assessing the quality of microbial genomes recovered from isolates, single cells, and metagenomes. Genome research 25, 1043–1055 (2015).

40. Orakov, A. et al. Gunc: detection of chimerism and contamination in prokaryotic genomes. Genome biology 22, 1–19 (2021).

41. Prasoodanan P.K. V. et al. A census of hidden and discoverable microbial diversity beyond genome-centric approaches. bioRxiv 2025.06.26.661807 (2025).

42. Aroney, S. T. N., Newell, R. J. P., Tyson, G. W. & Woodcroft, B. J. Bin chicken: targeted metagenomic coassembly for the efficient recovery of novel genomes. bioRxiv 2024.11.24.625082 (2024).

43. Schechter, M. S. et al. Quantitative insights into the efficacy of genome-resolved surveys of microbial communities through ribosomal protein phylogeography and EcoPhylo. bioRxivorg 2025.01.15.633187 (2025).

44. Quinlan, A. R. & Hall, I. M. BEDTools: a flexible suite of utilities for comparing genomic features. Bioinformatics 26, 841–842 (2010).

45. Meyer, F. et al. Amber: assessment of metagenome binners. GigaScience 7, giy069 (2018).

